# Intraspecific proteomic profiling and potential biological activities of the honey bee hemolymph

**DOI:** 10.1101/2023.02.15.528732

**Authors:** Salma A. Elfar, Iman M. Bahgat, Mohamed A. Shebl, Mathieu Lihoreau, Mohamed M. Tawfik

## Abstract

Pollinator declines have raised major concerns for the maintenance of biodiversity and food security, calling for a better understanding of environmental factors that affect their health. Here we used hemolymph analysis, a cheap, simple, yet powerful approach, to monitor the health state of Western honey bees *Apis mellifera*. We evaluated the intraspecific proteomic variations and the biological activities of hemolymph of bees collected from four Egyptian localities characterized by different food diversities and abundances. Lowest protein concentrations and the weakest bioactivities were recorded in hemolymph of bees artificially fed sucrose solution and no pollen. By contrast, highest protein concentrations and activities were recorded in bees that had the opportunity to feed on various natural resources. While future studies should expand comparisons to honey bee populations exposed to more different diets, our results strongly suggest hemolymph samples can be used as reliable indicators of bee nutrition and health.

## 1. Introduction

Bees are key pollinators [1,2] whose widespread declines have raised major concerns for the maintenance of terrestrial biodiversity and food security [3]. Over the past decades, the accelerated loss of bee populations has called for a better understanding of the environmental stressors that population growth and their mechanism of action, for instance through large-scale monitoring of bee health statuses across different habitats [4].

Like all insects, bees have an open circulatory system composed of hemolymph, which is an equivalent fluid to the blood in higher vertebrates [5]. Its main components are water, carbohydrates, proteins, inorganic salts, lipids, hormones and immune cells. As such hemolymph plays a very important and vital role to bees as it manages the distribution of nutrients along the body by supplying these nutrients to the tissues and organs [6]. Furthermore, it is the main site for defense against infections [7]. Indeed, the hemolymph of honey bees is well known to have antimicrobial, antioxidant and different immuno-modulatory agents [8,9]. The proteomic compositions of bee’s hemolymph can act as indicators to certain physiological and immune information that can monitor the health state of bees[5,10]. Therefore, hemolymph analysis can serve as simple, yet potentially powerful, means for monitoring bee population health status.

Variations in hemolymph composition is strongly linked to the nutrition of bees, since food is the major contributor to hemolymph proteins [11–13]. For instance, high hemolymph protein levels minimize the susceptibility of honey bees to pathogens [14,15]. By contrast, low hemolymph protein levels are a signature of poor health status. Previous research shows the hemolymph of the honey bee *Apis mellifera* varies in protein composition across individuals of the same population, especially within castes and among different developmental stages [16]. However, little is known concerning the variation in hemolymph protein composition among workers with consideration of diet variations. Moreover, the type of solvent used to extract hemolymph samples in the previous different studies affects solubility and structure of proteins, and thus the content and biological activities of hemolymph [17].

To address this question, we evaluated the intraspecific proteome variations and some biological activities of hemolymph of honey bee *A. mellifera* collected from different localities across Egypt. Egypt has both high biodiversity of bees and important differences of flower abundances and diversity across different regions. We therefore aimed to cover this variability by sampling bees from four locations characterized by contrasted diets (natural vs. artificial diets). We ran these analyses with the complementary solvents phosphate buffer saline and dimethyl sulfoxide, to make sure our analysis was as exhaustive as possible.

## 2. Materials and Methods

### 2.1. Chemicals

All chemicals used were purchased from Sigma Chemical Co. (St. Louis, MO). Phosphate buffer saline (PBS) (hydrophilic) or Dimethyl sulfoxide (DMSO) (hydrophobic) were used as solvents for the extraction of honey bee hemolymph samples. These two complementary solvents were used in order to extract as many molecules as possible.

### 2.2. Sample collection and preparation

Sweep nets were used to collect honey bee foragers in the different study sites (ca. 100 bees per location) from May to July 2021. Bees from the faculty of science of Port Said Governorate originated from colonies enclosed in a big flight tent and maintained in an artificial beehive with access to sugar syrup only. Bees from Ismailia, Suez governorates and Saint Catherine had access to different cultivated plants (see details in Table 1), thus providing diverse nutritional resources. The bees were kept at −20 °C until hemolymph collection.

**Table 1.** Localities and plantations of the study areas

| Symbol | Locality | Main feeding diet | Main plantation |
| --- | --- | --- | --- |
| A | Port said Governorate<br>31.259N 32.27E | Artificial (Sucrose syrup) | N.A |
| B | Ismailia Governorate<br>30.61N 32.25E | Natural | Cotton. |
| C | Suez Governorate<br>30.02N 32.34E | Natural | Corn, cucumber, okra and sesame. |
| D | Saint Catherine<br>28.56N 33.94E | Natural | Medicinal plants (thyme and sial tree) |

### 2.3. Hemolymph collection and preparation

Hemolymph was extracted by puncturing the coxal membrane using sterile insulin syringes and pressing the abdomen of bees. Clear and slightly yellow hemolymph was drawn out from the wound at the coxal area [15]. If cloudy yellow intestinal contents were taken, the sample was discarded. The hemolymph samples were kept into sterile Eppendorf tubes and stored at −20 °C until lyophilization. Equal weights of lyophilized hemolymph were dissolved in DMSO or PBS.

### 2.4. Protein Estimation

Protein concentration of each extract was measured in mg/ml by using Thermo Scientific™ Nano Drop™ One Micro volume UV-Vis Spectrophotometer, A280 for protein concentrations by using Bovine serum albumin (BSA) as a standard [18].

### 2.5. SDS-polyacrylamide gel electrophoresis

Protein gel profiles were obtained to investigate variations among samples by using Sodium dodecyl sulphate polyacrylamide gel electrophoresis (SDS-PAGE) for separating proteins depending on their molecular weights according to Laemmli [19] with some modifications. SDS-PAGE was performed at a total acrylamide concentration of 12%. The tested samples were mixed with the solubilizing buffer that contains 62.5 mM Tris-HCl (pH 6.8), 20% glycerol, 25(w/v) SDS, 0.5% 2-mercaptoethanol and 0.01% bromophenol blue, then heated for 4 min at 95°C. The samples were immediately loaded into wells, and electrophoresis was carried out at constant 35 mA for 2 h using Consort N.V. (Belgium) mini vertical electrophoresis system with running buffer. Finally, the gel was stained with 0.1% Coomassie Brilliant Blue (R-250) for the protein bands to be visualized.

### 2.6. HPLC (High performance liquid chromatography) analysis

Reversed Phase HPLC chromatograms of the honey bee hemolymph extracts separate the protein peaks based on their retention time. Each of the lyophilized hemolymph samples was dissolved in either solvent (PBS or DMSO) at the same concentration (5 mg/ml). Then, the samples were analysed using YL9100 HPLC System under the following conditions: C18 column (Promosil C18 Column 5 μm, 150 mm×4.6mm) as stationary phase and acetonitrile (ACN) gradients in the range of 10% to 100% acetonitrile in water for 50 min as mobile phase at flow rate 1 ml/min and by using UV detector at wavelength 280 nm [8].

### 2.7. Anticancer activity (MTT-assay)

The anticancer activities of the hemolymph extracts were experimented *in vitro* to assess activities and intraspecific variations among different hemolymph extracts when using PBS or DMSO. Human hepatocellular carcinoma (HepG2) and human cervical carcinoma (HeLa) cells were obtained from the Holding company for biological products and vaccines (VACSERA), Giza, Egypt. Cells were seeded in 96-well plate for 24 h (5×10^3^cells/well). After incubation, the cells were treated with serial concentrations of the hemolymph extracts in each solvent (1000, 500, 250, 125, 62.5, 31.25 μg/ml) and then incubated at 37 °C in 5% CO_2_ atmosphere for 48 h.

The cytotoxic effect of extracts on the proliferation of cells was evaluated using the 3-[4, 5-methylthiazol-2-yl]-2, 5-diphenyl-tetrazolium bromide (MTT) assay [20]. After 48 h treatment, medium including MTT dye was added to cells and incubated at 37°C for 4 h, to allow the production of formazan crystals in the viable cells only. DMSO was used in order to solubilize formazan crystals. Finally, the absorbance was measured at 540 nm using a Bio-Tek microplate reader ELISA. Each experiment was performed in triplicate, and the percentage of cell viability was calculated by using the following equation:

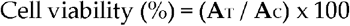

where: **A**_T_: absorbance of treated cells, **A**_C_: absorbance of the control cells (untreated cells).

The concentration that inhibits the growth of 50% of the cells (IC_50_) values was determined for each sample from a dose-response curve between dose concentration (X-axis) and cell inhibition percentage (Y-axis).

### 2.8. Antimicrobial activity screening

The antimicrobial activity of hemolymph extracts was experimented to evaluate the activity variations against bacteria by using agar well diffusion method [21]. Antimicrobial activity was determined against two Gram positive bacteria (*Bacillus subtilis and Staphylococcus aureus*) and two Gram negative bacteria (*Escherichia coli and Salmonella typhimurium*). Briefly, the agar plate surface was inoculated by spreading a volume of the microbial inoculum over the entire agar surface, then making a hole with a diameter of 6 to 8 mm aseptically by using a sterile tip. A volume of about (20-100μL) of each extract or control at known concentrations was introduced into the well. After that, the agar plates were incubated at suitable conditions depending on the microorganism used for the test. Gentamycin was used as a positive control while PBS and DMSO were used as negative controls. The antibacterial activities of the extracts were expressed as inhibition zones in millimetres (mm) [22,23].

### 2.9. DPPH radical scavenging assay

The antioxidant ability of the hemolymph extracts was experimented in vitro to demonstrate the variations among the extracts. The radical scavenging capacity of the investigated hemolymph extracts on the stable free radical 1,1-Diphenyl-2-picrylhydrazyl (DPPH) was assessed according to the method of Braca [24] with some modifications. Briefly, 100μl of prepared DPPH methanolic solution (0.004% in methanol) with 0.3 ml of sample extracts or standard and incubated in dark for 30-60 min at 25 °C. Ascorbic acid (Vitamin C) was used as a standard (positive control), and the absorbance was measured at 515 nm by using spectrophotometer. Negative control was prepared by adding 2.7 mL of DPPH in 0.3 mL of the solvent used in the extract (AC). The antioxidant activity of the samples was calculated from the following equation [25,26]:

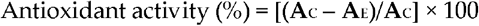

where: **A**_C_: The mean of absorbance of negative control and **A**_E_: The mean of absorbance of extract or standard.

### 2.10. Hemolytic activity assay

The ability of the hemolymph extracts to lyse erythrocytes was evaluated to demonstrate the selectivity and safety of hemolymph for future applications in experimental animal models regarding preclinical medical research approaches The hemolytic activity of the bee’s hemolymph was evaluated on human erythrocytes [27]. Fresh blood samples were collected in test tubes containing anticoagulant (EDTA) then centrifuged for 5 min at 2000 rpm. Blood samples were washed several times with sterile PBS then series of various concentrations of the extracts were added to RBCs and incubated for an hour at room temperature. After incubation, the samples were centrifuged for 5 min at 10000 rpm and then the absorbance of the released hemoglobin was measured at 570 nm. 10% Triton X-100 was used as a positive control (100% hemolysis) and sterile PBS and DMSO were used as negative controls (0% hemolysis). This experiment was carried out three times and the hemolysis percentage was calculated for each sample by using the equation:

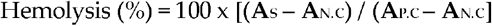

where: **A**_S_: the absorbance of the sample, **A**_N.C_: the absorbance of the negative control, **A**_P.C_: is the absorbance of the positive control.

### 2.11. Statistical analysis

Statistical analysis was done using SPSS 22.0. Data were analysed using Student’s t-tests or One-way ANOVAs followed by a Tukey’s test. When the P-value was lower than 0.05, the difference in the means of the samples were considered as statistically significant. Means are reported with standard errors (mean ± SE).

## 3. Results

### 3.1. Naturally fed honey bees had higher potein concentration in hemolymph

The amount of proteins found in each hemolymph extract was measured spectrophotometry (**Figure 1**). Among the DMSO dissolved samples, extracts from site **B** had higher protein concentrations (0.12 ± 0.0006 mg/ml) than extracts from all other sites. Among the PBS dissolved samples, extracts from site **D** had the highest protein concentration (0.19 ± 0.03mg/ml). Extracts from site **A** had a significantly lower protein concentration than extracts from all the other sites in either PBS or DMSO (0.006 ± 0.0005 mg/ml and 0.02 ±0.0003 mg/ml) respectively. These results showed variations in the protein concentrations among the bee hemolymph extracts with different feeding diets. Lowest protein concentrations were consistently recorded for the artificially fed bees in site **A**.

**Figure 1.**
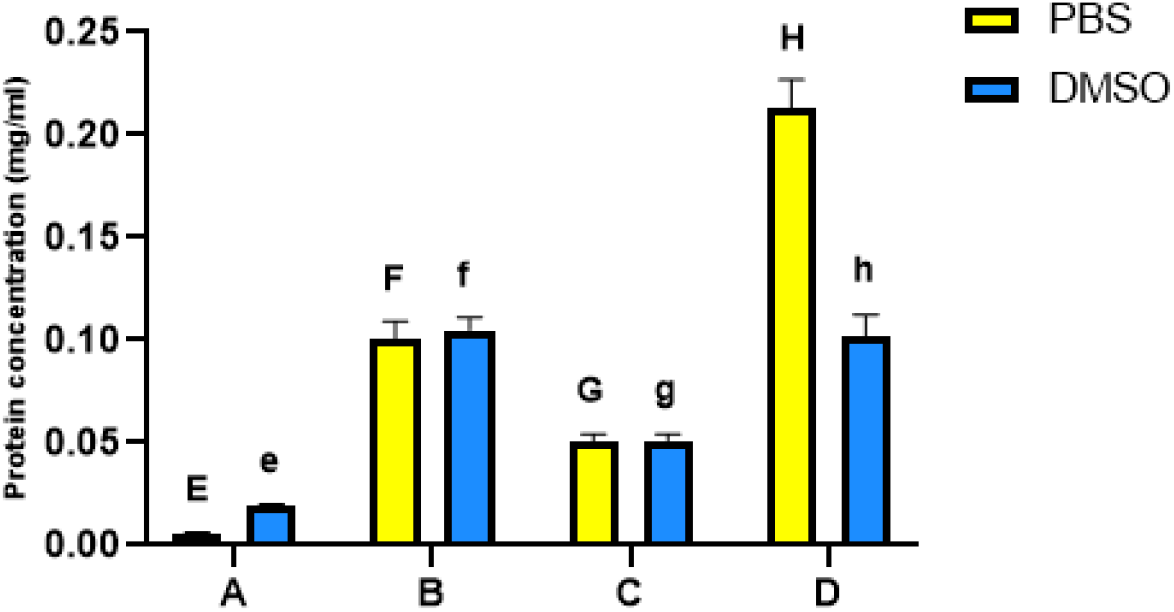
Protein concentrations in mg/ml for honey bee hemolymph extracted in either solvents PBS or DMSO where, (**A**): Bee’s hemolymph from a hive in Port Said, (**B**): from Ismailia governorate, (**C**): from Suez governorate, (**D**): from Saint Catherine. Bars and error bars represent the mean values ±SE obtained from triplicate measurments. Different letters above bars indicate significant differences (Tukey’s test, P < 0.05): uppercase letters show comparisons across PBS dissolved samples; lowercase letters show comparisons across DMSO dissolved samples.

### 3.2. Honey bee hemolymph showed different protein composition across sites

Quantitative analysis of proteins in the hemolymph extracts revealed a variety of protein bands ranging in mass from 5 to ~250 kDa when using either PBS or DMSO solvents (**Figure 2**).

**Figure 2.**
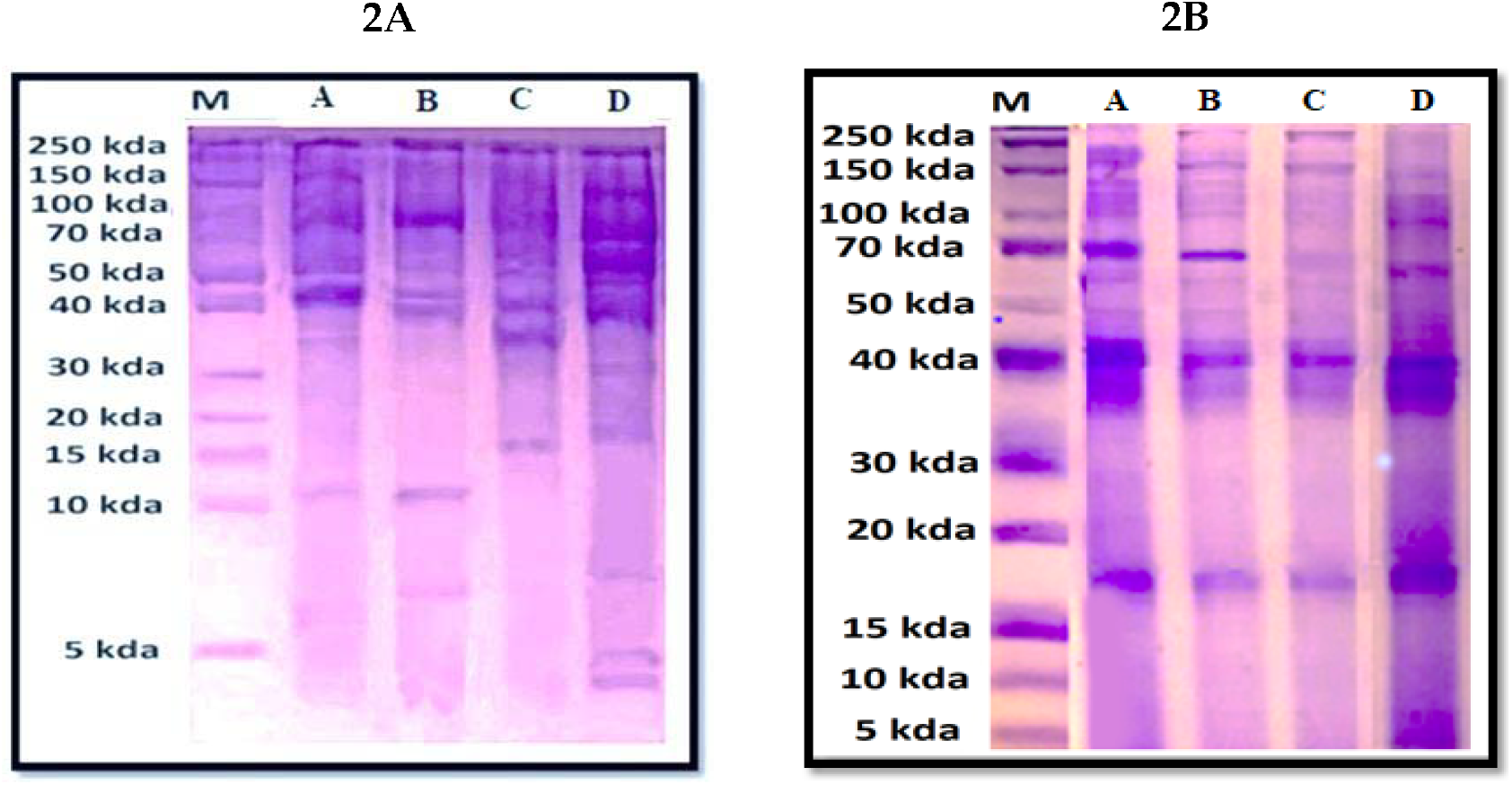
SDS-PAGE gel for honey bee hemolymph extracts dissolved in PBS (Figure 2A) or DMSO (Figure 2B) where, (**A**): Bee’s hemolymph from a hive in Port Said, (**B**): from Ismailia governorate, (**C**): from Suez governorate, (**D**): from Saint Catherine. **M**: marker. The molecular weights of the proteins and their amounts are expressed by colour intensity of the protein bands.

The SDS profile of the samples dissolved in PBS (**Figure 2A**) revealed six common bands with molecular weights ~40, ~50, ~60, ~100, ~180 and ~250 kDa. A band with molecular weight ~10 kDa was only recorded in extracts from sites A and B. A band with molecular weight ~17 kDa was only recorded in extracts from sites C and D. Three unique bands with molecular weights (~5, ~30 and ~70 kDa) were specific for (D) extracts.

Among the samples dissolved in DMSO (**Figure 2B**), the common bands were observed at molecular weights ranging from ~17 to ~250 kDa. A unique band with molecular weight ~5 kDa was recorded only in extracts from site D.

HPLC analysis was then used to separate the proteins of the hemolymph extracts according to their peaks expressed in retention time. The RP-HPLC chromatograms of the hemolymph extracts dissolved in PBS revealed five common peaks in extracts from sites B, C and D at retention times (7.6, 12.4, 19.9, 25.6 and 43.0 minutes). Another five common peaks were recorded at retention times (22.1, 28.7, 30.4, 33.2 and 38.7 minutes) among the extracts from all sites when using DMSO as a solvent (Table 2). Chromatograms are supplemented in the supplementary Figures A1 and A2.

**Table 2.** RP-HPLC chromatogram peaks profiles of the tested honey bee hemolymph extracted in PBS or DMSO.

| Retention time (minutes) | Peak area (x10<sup>2</sup> mV.s) |  |  |  |  |  |  |  |
| --- | --- | --- | --- | --- | --- | --- | --- | --- |
|  | A |  | B |  | C |  | D |  |
|  | PBS | DMSO | PBS | DMSO | PBS | DMSO | PBS | DMSO |
| 2.9 | N.A | N.A | 1.8 | N.A | 12.3 | N.A | N.A | N.A |
| 4.0 | N.A | N.A | N.A | 27.7 | N.A | 1167 | N.A | 0.2 |
| 4.3 | N.A | N.A | N.A | N.A | 14.4 | N.A | 3386 | N.A |
| 5.0 | N.A | N.A | N.A | 838.7 | N.A | N.A | N.A | 397 |
| 6.7 | 36.4 | N.A | 3.9 | N.A | 1.0 | N.A | N.A | N.A |
| 7.6 | N.A | N.A | 4.6 | N.A | 0.8 | N.A | 163.5 | N.A |
| 10.3 | N.A | N.A | N.A | N.A | 4.8 | N.A | 247.5 | N.A |
| 11.3 | N.A | N.A | 2.3 | N.A | 3.9 | N.A | N.A | N.A |
| 12.4 | N.A | N.A | 10.9 | N.A | 2.5 | N.A | 1265 | N.A |
| 14.4 | N.A | N.A | 5.9 | N.A | N.A | N.A | 105.3 | N.A |
| 19.9 | N.A | N.A | 4.1 | N.A | 5.0 | N.A | 363.7 | N.A |
| 22.1 | N.A | 0.3 | N.A | 1.5 | N.A | 0.6 | N.A | 2.6 |
| 23.2 | N.A | N.A | N.A | 3.6 | N.A | 1.0 | N.A | N.A |
| 23.5 | N.A | 0.9 | N.A | N.A | N.A | N.A | N.A | 1.2 |
| 25.6 | N.A | N.A | 9.1 | N.A | 18.0 | N.A | 770.9 | N.A |
| 28.7 | N.A | 0.6 | N.A | 2.6 | N.A | 0.3 | N.A | 1.8 |
| 28.8 | 43.2 | N.A | 2.3 | N.A | 4.7 | N.A | N.A | N.A |
| 30.3 | 94.8 | N.A | 1.8 | N.A | 5.2 | N.A | N.A | N.A |
| 30.4 | N.A | 1.3 | N.A | 0.5 | N.A | 0.2 | N.A | 0.4 |
| 33.2 | N.A | 0.2 | N.A | 0.3 | N.A | 0.3 | N.A | 0.2 |
| 34.8 | N.A | N.A | N.A | 0.8 | N.A | N.A | N.A | 1.4 |
| 36.5 | N.A | 1.0 | N.A | 0.7 | N.A | N.A | N.A | 0.2 |
| 38.1 | N.A | N.A | 9.1 | N.A | 3.0 | N.A | N.A | N.A |
| 38.7 | N.A | 10.5 | N.A | 0.2 | N.A | 0.1 | N.A | 0.1 |
| 40.7 | N.A | N.A | N.A | 0.1 | N.A | N.A | N.A | 0.1 |
| 43.0 | N.A | N.A | 4.4 | N.A | 2.6 | N.A | 99.1 | N.A |
| 47.7 | N.A | N.A | 14.6 | N.A | 10.7 | N.A | N.A | N.A |

### 3.3. Honey bee hemolymph suppresses the growth of HepG2 and HeLa cells

The anticancer activity of the hemolymph extracts were analysed in vitro. The cytotoxic effects of honey bee hemolymph samples were evaluated against the viability of HepG2 and HeLa. The viability of the two cancer cell lines was inhibited in a dose-dependent manner after being treated with different concentrations of the hemolymph extracts in either PBS or DMSO after 48 h of treatment (**Figure 3**). Overall, DMSO dissolved extracts possessed higher anti-proliferative activity than those dissolved in PBS (**Figure 4**).

**Figure 3.**
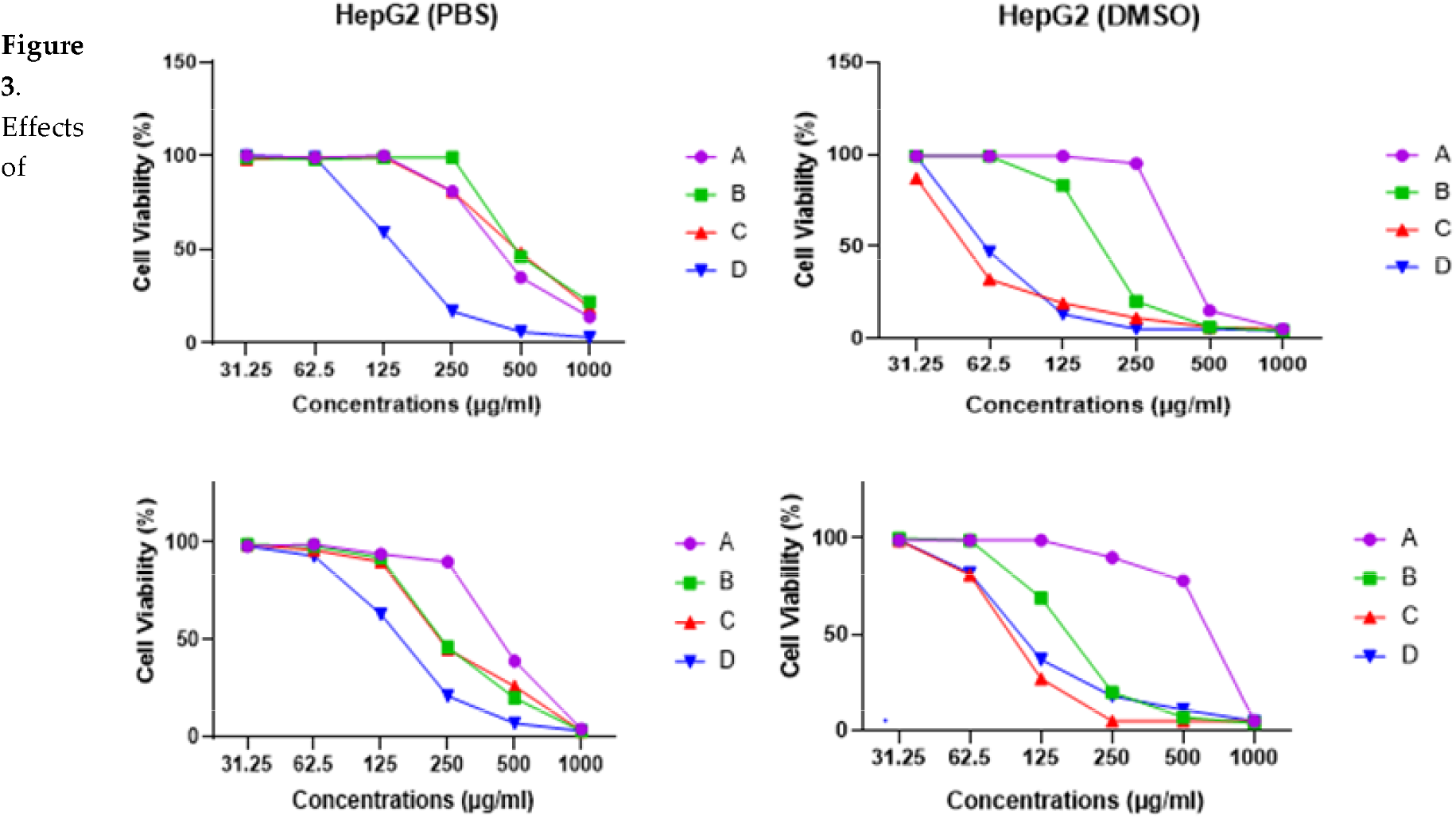
Effects of hemolymph extracts of honey bees on human cancer cell proliferation (HepG2 and HeLa) at different concentrations in either solvents PBS or DMSO where, (A): Bee’s hemolymph from a hive in Port Said, (B): from Ismailia governorate, (C): from Suez governorate, (D): from Saint Catherine.

**Figure 4.**
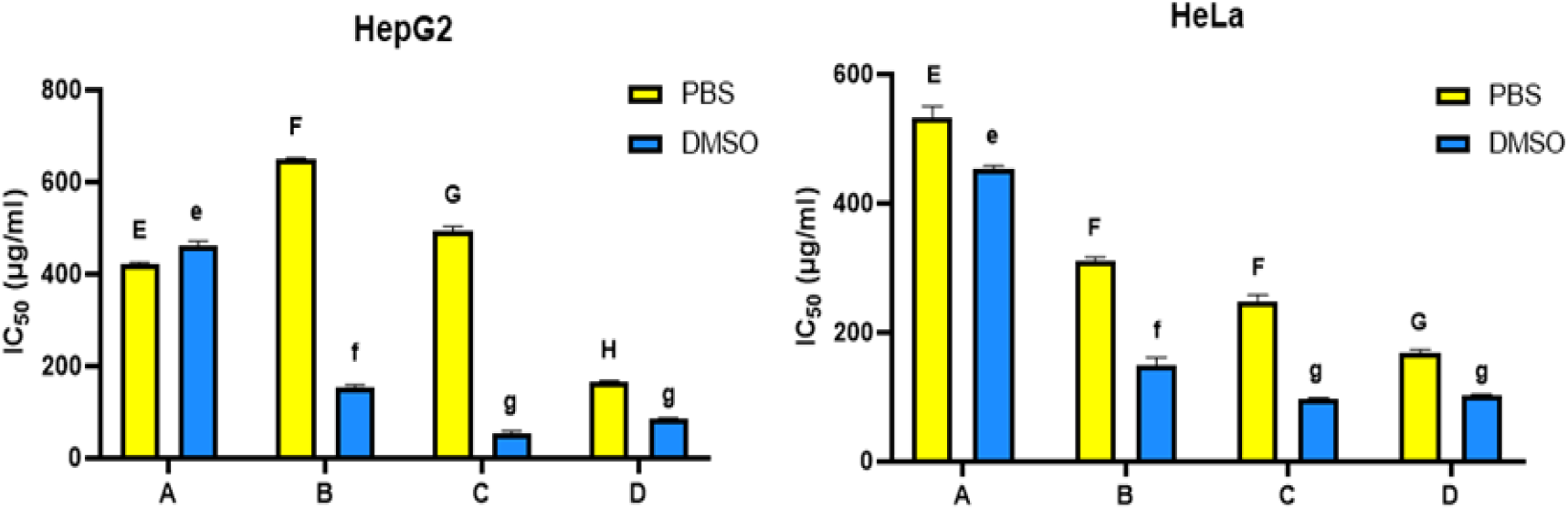
IC_50_ in microgram/ml of hemolymph extracts of honey bees against Hepg2 and HeLa cell lines, in either solvents PBS or DMSO where, (**A**): Bee’s hemolymph from a hive in Port Said, (**B**): from Ismailia governorate, (**C**): from Suez governorate, (**D**): from Saint Catherine. Bars and error bars represent the mean values ± SE obtained from triplicate measurments. Different letters above bars indicate significant differences (Tukey’s test, P < 0.05): uppercase letters show comparisons across PBS dissolved samples; lowercase letters show comparisons across DMSO dissolved samples.

Extracts from site C dissolved in DMSO showed the highest cytotoxic effects against HepG2 and HeLa cells (IC_50_= 52.03 & 97.95 μg/ml respectively). The lowest cytotoxic effect was recorded for extracts from site A (IC_50_= 463.36 & 454.02μg/ml) (**Figure 4**).

When using PBS as a solvent, higher cytotoxic effect was recorded for extracts from site D (IC_50_= 164.5 & 169.7 μg/ml against HepG2 and HeLa respectively). On the other hand, extracts from sites A and B showed the lowest cytotoxic effects against HeLa (IC_50_= 533.08μg/ml) and HepG2 (IC_50_= 649.36μg/ml) (**Figure 4**).

### 3.4. Honey bee hemolymph inhibts the growth of Gram positive and Gram negative bacteria

The antimicrobial activity of the hemolymph extracts from sites A, B, C and D in either solvent (PBS or DMSO) was analysed against two Gram positive bacteria and two Gram negative bacteria using well diffusion method (**Table 3**). Overall, PBS samples revealed higher antibacterial activity than DMSO samples (P<0.05). By using either PBS or DMSO solvents (**Table 3**), the highest antibacterial activities were observed for the extracts from site B against *E. coli* with inhibition zones (40 mm) and (38 mm) respectively. However, extracts from site A showed the lowest antibacterial activities against all tested bacteria with inhibition zone ranging from 19 to 28 mm (**Figure 5**).

**Table 3.** Inhibition zone (mm) of honey bee hemolymph extracted in either solvents PBS or DMSO on various types of bacteria (*Bacillus subtilis*, Staphylococcus *aureus, Escherichia coli, Salmonella typhimurium*).

| Sample<br>Pathogen | PBS |  |  |  | DMSO |  |  |  | CON. |
| --- | --- | --- | --- | --- | --- | --- | --- | --- | --- |
|  | A | B | C | D | A | B | C | D |  |
| *Bacillus subtilis* | 28±1.2<sup>a</sup> | 34±1.0<sup>b</sup> | 30±0.9<sup>a</sup> | 27±0.9<sup>a</sup> | 26±0.6<sup>a</sup> | 32±1.6<sup>b</sup> | 31±1.5<sup>ab</sup> | 25±1.0<sup>ac</sup> | 22±0.6 |
| *Staphylococcus aureus* | 23±1.0 | 25±2.0 | 28±0.9 | 28±1.0 | 21±1.6 | 24±0.9 | 23±1.0 | 25±1.9 | 15±0.9 |
| *Escherichia coli* | 21±1.3<sup>a</sup> | 40±1.5<sup>b</sup> | 32±0.5<sup>c</sup> | 25±1.3<sup>a</sup> | 19±1.3<sup>a</sup> | 38±1.0<sup>b</sup> | 27±1.7<sup>c</sup> | 23±0.7<sup>ac</sup> | 17±1.0 |
| *Salmonella typhimurium* | 22±1.3 | N.A | 26±1.3 | N.A | 20±1.5 | N.A | 25±2.2 | N.A | 23±0.7 |

Where **NA** = no activity, (**A**): Bee's hemolymph from a hive in Port Said, (**B**): from Ismailia governorate, (**C**): from Suez governorate, (**D**): from Saint Catherine, **CON**: Gentamycin as positive control. Mean values ± SE were obtained from triplicate measurements. The values with different superscript letters (a–d) in a rows are significantly different across locations (Tukey test, $P<0.05$ ).

**Figure 5.**
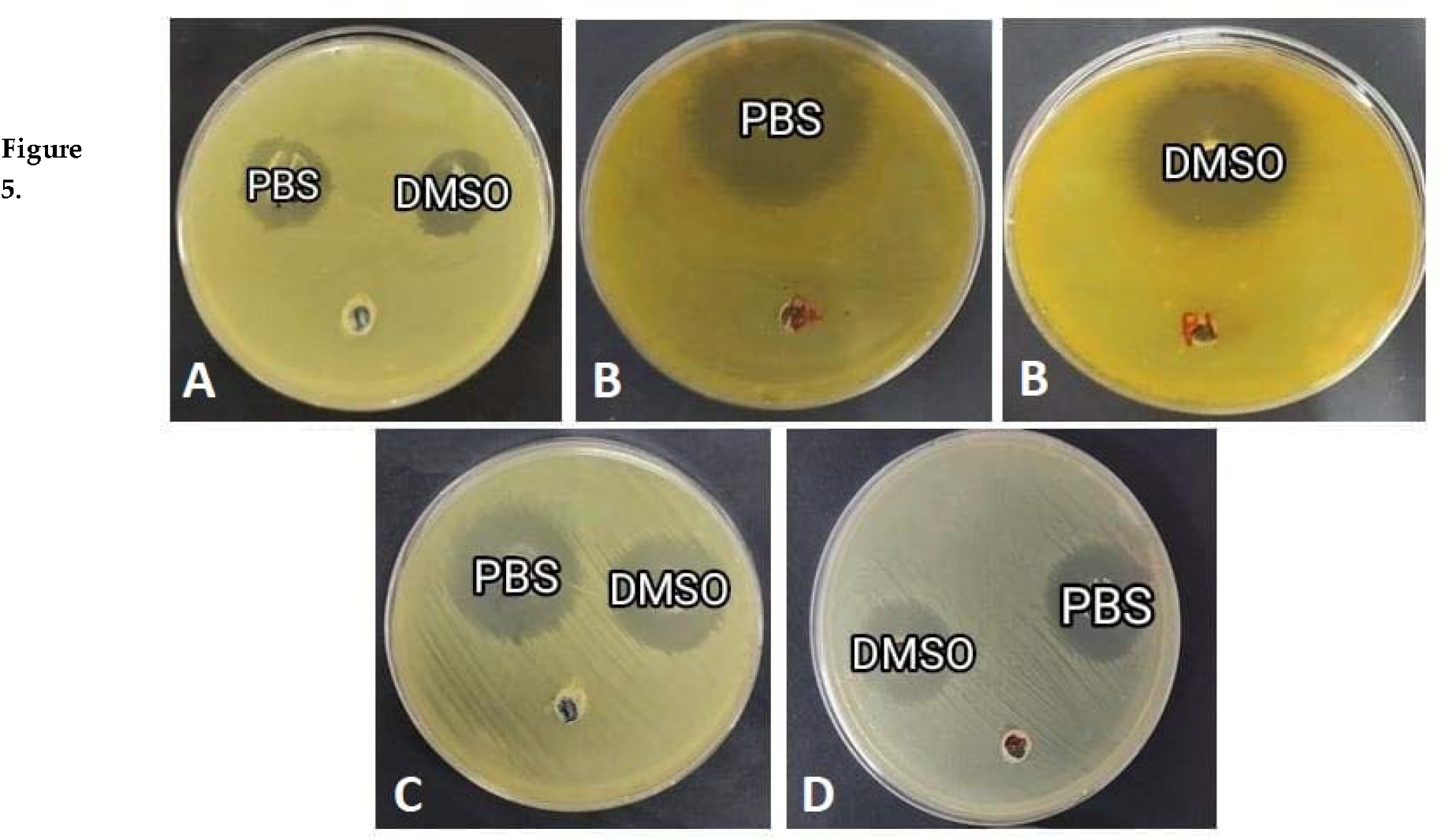
Antimicrobial activity of honey bee hemolymph extracted in either solvents PBS or DMSO against *E.coli*, where, (**A**): Bee’s hemolymph from a hive in Port Said, (**B**): from Ismailia governorate, (**C**): from Suez governorate, (**D**): from Saint Catherine.

### 3.5. Honey bee hemolymph scavenges DPPH free radical

The antioxidative activity of the hemolymph extracts on DPPH free radicals showed a potent radical scavenging activity in either PBS or DMSO solvents. Overall, hemolymph extracts in PBS solvent displayed higher antioxidant properties than that of hemolymph extracts in DMSO solvent (Figure 6) (P<0.05).

**Figure 6.**
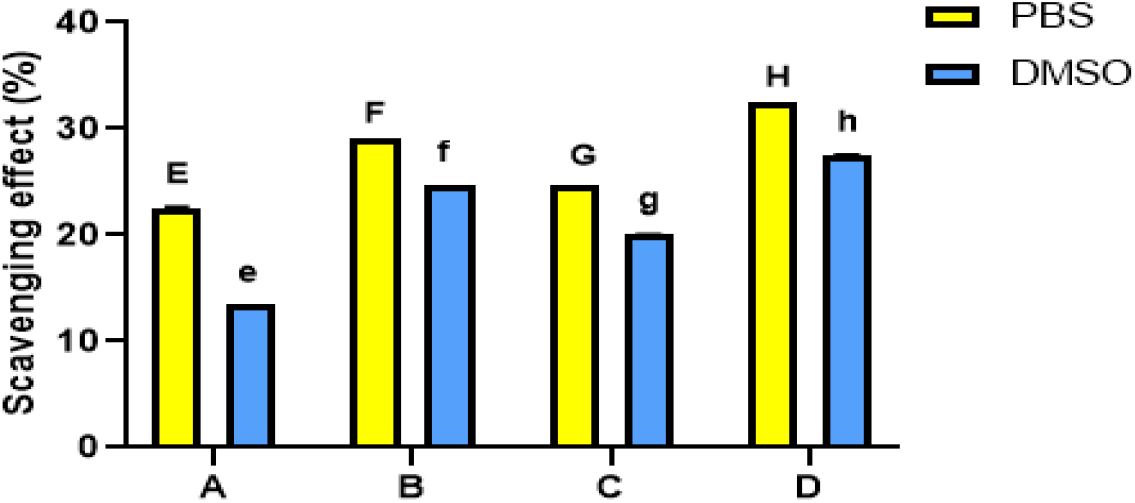
DPPH radical scavenging activity of honey bee hemolymph extracted in either solvents PBS or DMSO where, (**A**): Bee’s hemolymph from a hive in Port Said, (**B**): from Ismailia governorate, (**C**): from Suez governorate, (**D**): from Saint Catherine, bars and error bars represent the mean values ±SEM obtained from triplicate measurments. Bars and error bars represent the mean values ± SE obtained from triplicate measurments. Different letters above bars indicate significant differences (Tukey’s test, P < 0.05): uppercase letters show comparisons across PBS dissolved samples; lowercase letters show comparisons across DMSO dissolved samples.

There was significant difference between the extracts in either PBS or DMSO solvents (P<0.001). Extracts from site D possessed the highest antioxidant activity in either of the two solvents (PBS or DMSO) (P<0.001). On the other hand, extracts from site A possessed the lowest scavenging activity (**Figure 6**).

### 3.6. Honey bees possess low hemolytic activity against human erythrocytes

Hemolytic activities of the hemolymph from honey bee extracts were evaluated on human erythrocytes. None of the honey bee extracts displayed hemolytic activity against erythrocytes when using either PBS or DMSO solvents at different concentrations (μg/ml) when compared to negative and positive controls (**Figure 7**).

**Figure 7.**
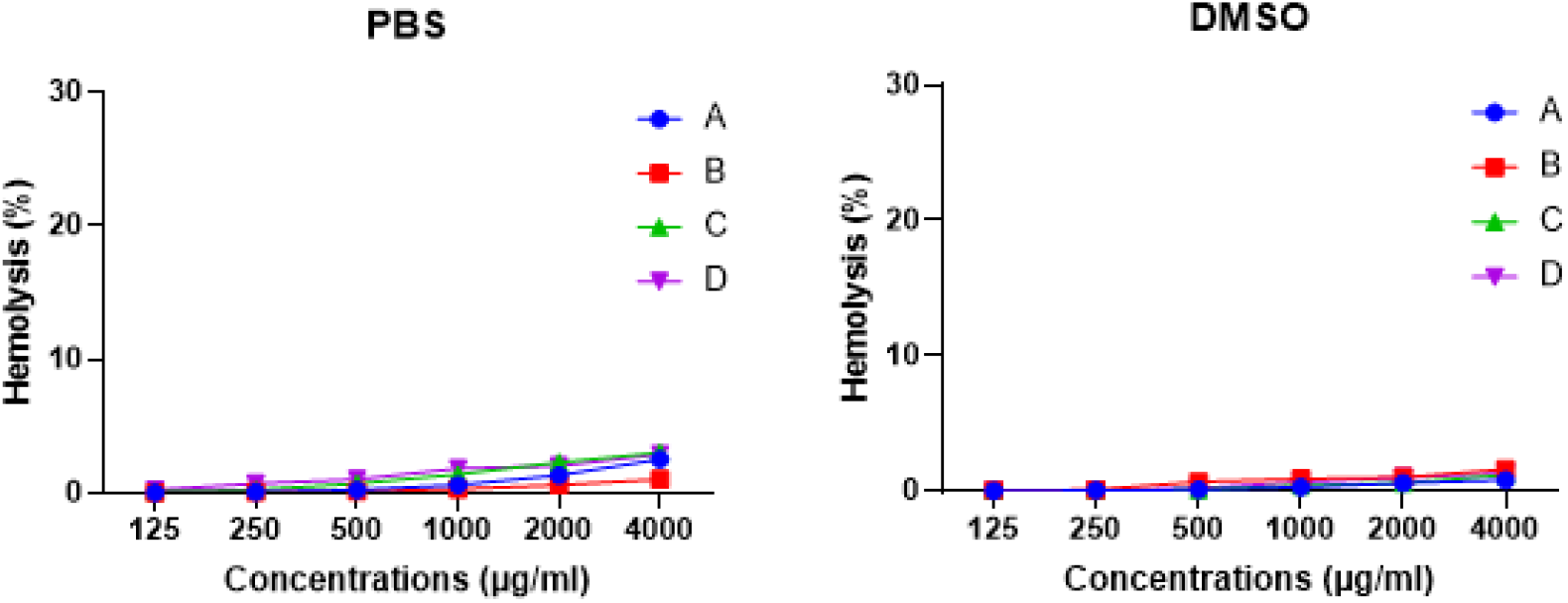
Hemolytic activity of honey bee hemolymph extracted in either solvents PBS or DMSO at different concentrations against erythrocytes where, (**A**): Bee’s hemolymph from a hive in Port Said, (**B**): from Ismailia governorate, (**C**): from Suez governorate, (**D**): from Saint Catherine.

## 4. Discussion

Our study describes intraspecific variations in the protein profiles and concentrations of the hemolymph of honey bees (*A. mellifera*) collected from different localities in Egypt chosen to offer contrasted diets to bees. Lowest protein concentrations and the weakest bioactivities were recorded in hemolymph of bees fed sucrose solution and no pollen. By contrast, high protein concentrations and activities were systematically recorded in bees that had the opportunity to feed on various natural resources

High protein concentrations were recorded in hemolymph of bees fed on natural resources as compared to bees fed sucrose solution. This is consistent with a study that reported the negative impact of sugar solution feeding diet on protein concentration of honey bees hemolymph [28]. Other previous studies have reported the influence of different feeding diets of honey bees on their hemolymph structure [29,30]. The protein content of hemolymph can be used to assess the efficacy of protein diets and pollen quality [11,31–33]. Our study shows it can also be used as an indicator of bee health since high protein concentrations of the hemolymph extend bee life span, improve immunity, and resistance to pathogens [34–36].

Our analysis also reveals important variations in protein concentration depending on the solvent used in extraction. PBS and DMSO were used to extract the honey bee hemolymph. PBS acts as a solvent for water soluble (hydrophilic) proteins [37], while DMSO acts as a solvent for lipid soluble (hydrophobic) proteins [38]. Proteins structure, stability and solubility are markedly affected by the polarity of solvent [17]. Consequently, the use of each of the two solvents explains variations in hemolymph biological activities, which may be related to their influence on hemolymph proteomes. We recommend future detailed analyses of hemolymph protein contents must therefore use these two complementary solvents for extraction.

The SDS-PAGE gel of the hemolymph of our extracts showed variations in their protein bands. In general, more protein bands were recorded when using DMSO than when using PBS. The protein bands with molecular weight (□17 kDa) were detected in most extracts dissolved in either PBS or DMSO. These bands are in the same range of a class of lectins, which is a common group of proteins in arthropods [39]. Insect lectins have several important roles in immune response, homeostasis and binding for the storage or transport of carbohydrates [40,41]. Lectins may also possess antimicrobial, antioxidant and anticancer activities and can be considered as a potential natural bioactive agent [42–46]. The protein bands with molecular weight (□60 kDa) were also found in all the tested PBS and DMSO dissolved extracts. Similar antimicrobial proteins have been fractionated from cockroach hemolymph [8].

A hemocyanin-like protein band (□70 kDa) was detected in all DMSO dissolved extracts and some of those dissolved in PBS. Hemocyanin is an oxygen transport protein that has a wide range of biological activities including antioxidant, anti-parasitic, antimicrobial and anticancer activities [45,47,48]. The protein bands with molecular weight (□110 kDa) were found in all PBS and DMSO dissolved extracts. These bands are similar to hexamerins, which are storage proteins that are derived from hemocyanin [49]. The protein bands with molecular weight (□180 kDa) were detected in all PBS and DMSO dissolved extracts. These bands are in the same range of vitellongenin in honey bee hemolymph [16]. Vitellogenin is a large female specific gluco-lipoprotein that can act as female nutrient storage and also lipid carrier protein [50,51]. Honey bee vitellogenin was shown to have antimicrobial, antioxidant and immunological activities [52].

Protein bands with molecular weight >250 kDa have been observed in the proteomic profiles of all PBS and DMSO hemolymph extracts. These bands correspond to apolipophorin, a lipid transporter lipoprotein [53]. These findings agreed with [54] and [16] who revealed similar bands in honey bee hemolymph.

Interestingly, chromatograms of honey bees hemolymph collected from Saint Catherine showed the highest peak areas, which is an indicator for the amount of proteins. This reinforces the results of the SDS-PAGE gel which reveals that the protein bands of extract from this location had the highest protein concentrations based on the colour intensity of the bands. Saint Catherine is the most floristically diverse regions in Middle East. It is an Egyptian natural protected area accounts for nearly one-fourth of the total flora of Egypt. Such vegetation diversity let bees in this region to have a foraging advantage which may probably enrich the protein content of honey bee hemolymph [55–58].

Both PBS and DMSO hemolymph dissolved extracts revealed a relative degree of cytotoxicity against HepG2 and HeLa in a dose dependent manner with preference for DMSO dissolved extracts. There was a variation in the cytotoxicity between the hemolymph samples in the two solvents against the two tested cell lines. This variation may be related to the significant differences in their protein profiles and antioxidant activities. The higher cytotoxicity of DMSO dissolved extracts against both cell lines may be attributed to the role of DMSO in dissolving lipoproteins [38]. Previous showed the effect of using lipoprotein complexes to improve the selectivity of anticancer drugs [59].

The diversity and prevalence of the bioactive proteins such as lectins, hemocyanin, vitellogenin, hexamerins and apolipophorin in honey bee’s hemolymph extracted either in PBS or DMSO may elucidate the potent biological activities of hemolymph in the present study. The results revealed that honey bee hemolymph extracts showed prominent antimicrobial activities against different species of Gram positive and Gram negative bacteria. *A. mellifera*, like other invertebrates, possesses a very effective innate immune mechanism to defend against microorganisms and pathogens [60]. The deficiency in the protein compositions of hemolymph affects the ability of the honey bee to resist diseases [61]. So, feeding on natural resources would have a great impact on honey bees hemolymph bioactivities [62,63].

All the tested hemolymph extracts dissolved in either PBS or DMSO solvents showed no hemolytic activity when experimented against human erythrocytes at different tested concentrations. This finding agreed with [64] who reported the non-hemolytic activity of honey bee hemolymph. Interestingly, in this latter study they demonstrate the potential anticancer effect of hemolymph mixed with herbals in mice models. In this regard, honey bee hemolymph may represent a natural potential selective therapeutic agent in cancer treatment.

## 5. Conclusions

Overall, our study highlights significant intraspecific variability in the composition and concentration of hemolymph of honey bees sampled in Egypt. The hemolymph of the honey bee that could feed on cultivated plants and wild medicinal plants, as in bees from Saint Catherine, possessed relatively higher protein concentrations compared to that of bees reared only on sugar solutions in artificial hives. While future studies should expand comparisons of bees exposed to more diets in more different environments, our results strongly suggest hemolymph samples can be used as simple, yet powerful, indicator of bee nutrition, health, and ultimately, environment quality to sustain populations of pollinators.

## Author Contributions

The study was planned and designed by MT, IB, MS, ML; field collection, laboratory studies and data analysis were done by SE, MT, IB, MS. All authors contributed in the writing of the manuscript. All authors have read and agreed with this version of the manuscript.

## Data Availability Statement

Data available on request due to restrictions, e.g., privacy or ethical.

## Acknowledgments

We are so grateful to Prof. Soliman Kamel and Mr. Saied Aboud for contribution in honey bee collection.

Conflicts of Interest: The authors declare no conflict of interest.

## Disclaimer/Publisher’s Note

The statements, opinions and data contained in all publications are solely those of the individual author(s) and contributor(s) and not of MDPI and/or the editor(s). MDPI and/or the editor(s) disclaim responsibility for any injury to people or property resulting from any ideas, methods, instructions or products referred to in the content.

## Supplementary materials

**Figure A1.**
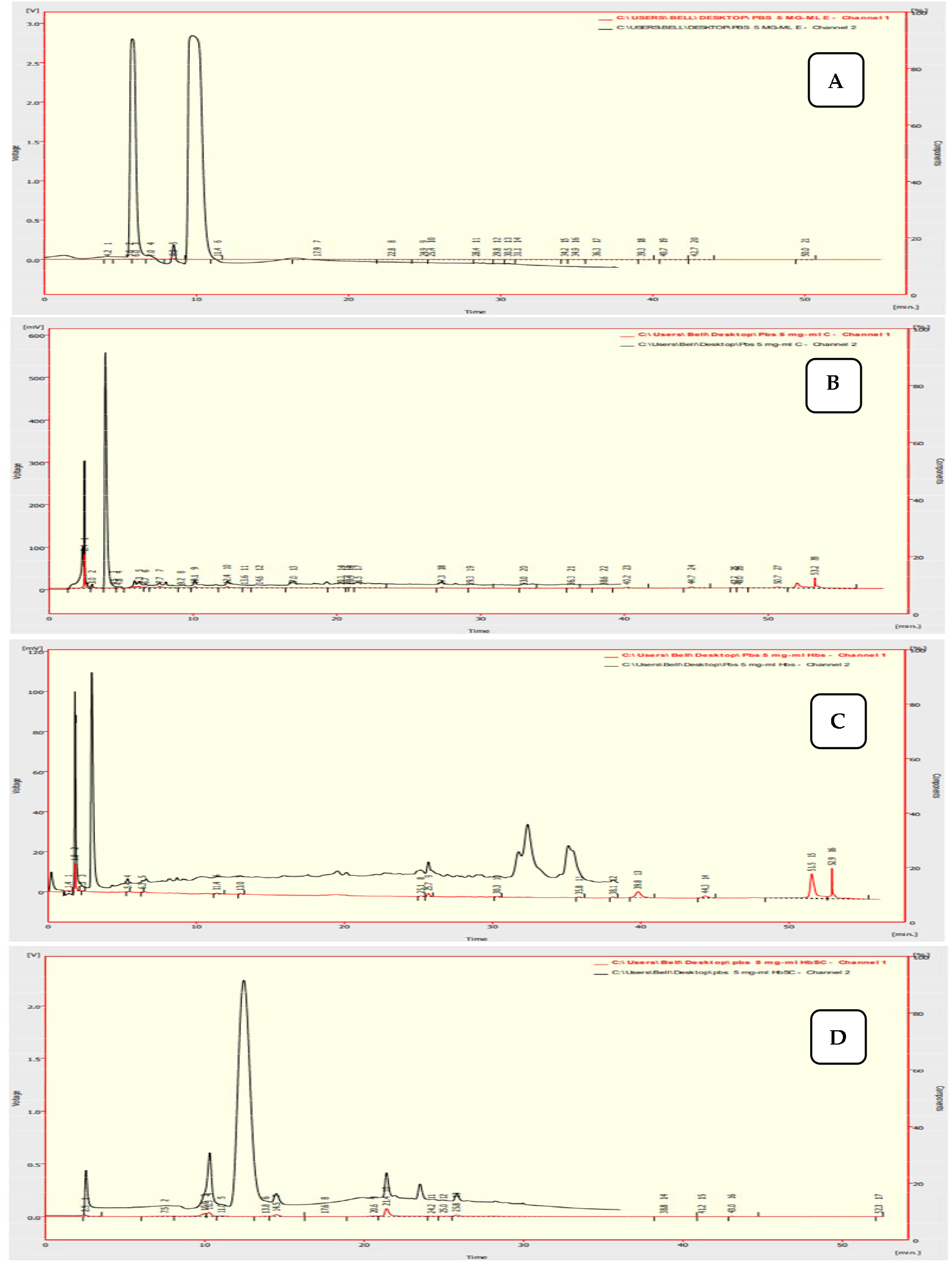
Chromatograms of RP-HPLC of the PBS dissolved honey bee hemolymph extracts where, (**A**): Bee’s hemolymph from a hive in Port Said, (**B**): from Ismailia governorate, (**C**): from Suez governorate, (**D**): from Saint Catherine

**Figure A2.**
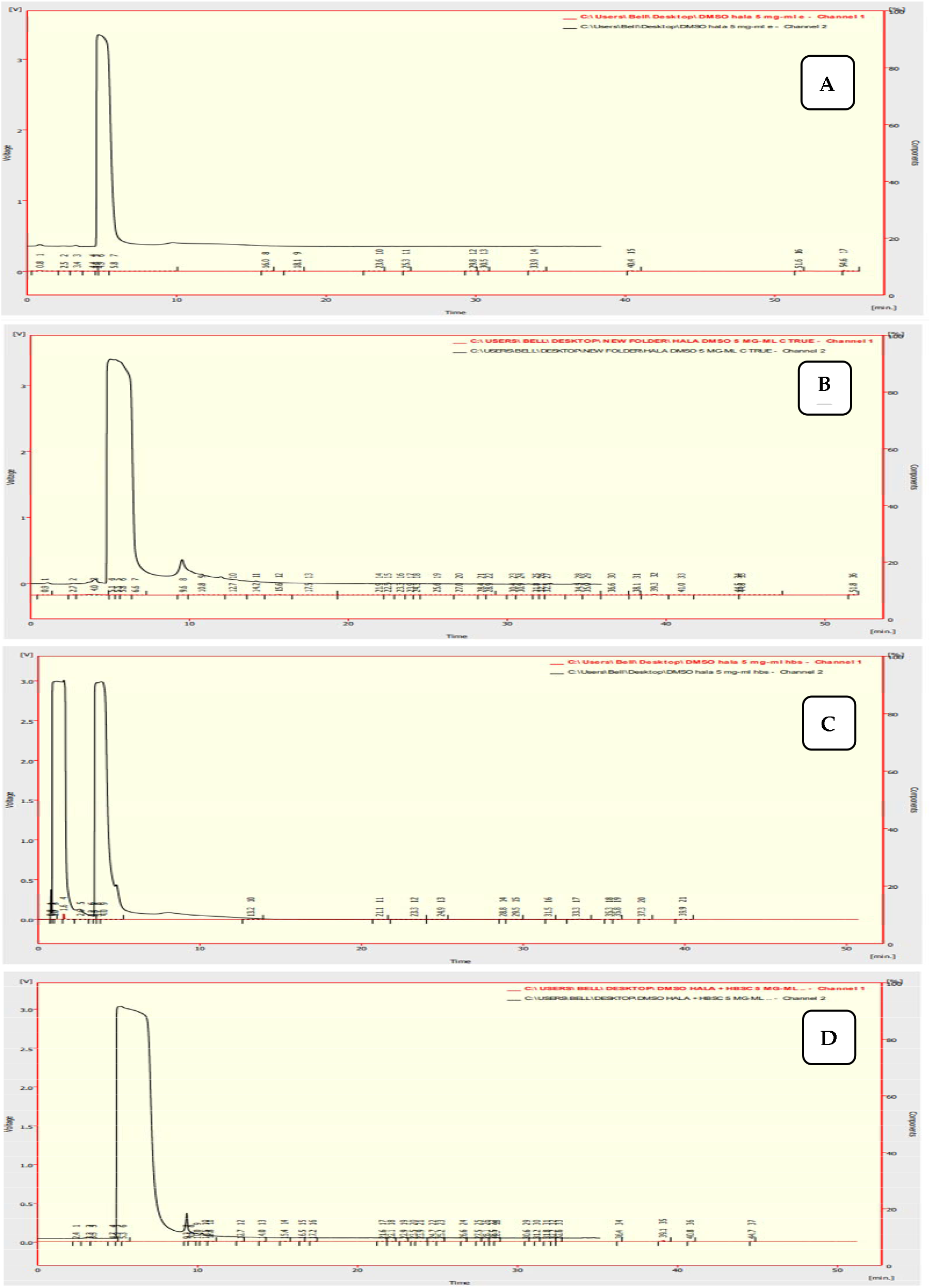
Chromatograms of RP-HPLC of the DMSO dissolved honey bee hemolymph extracts where, (**A**): Bee’s hemolymph from a hive in Port Said, (**B**): from Ismailia governorate, (**C**): from Suez governorate, (**D**): from Saint Catherine

